# Amoxicillin induces gut dysbiosis leading to long term suppression of type-17 immune tone in the lungs

**DOI:** 10.64898/2026.03.05.709937

**Authors:** Marika Orlov, Mallory Karr, Naoko Hara, James Needell, Carol M. Aherne, Jennifer L. Matsuda, Brent E. Palmer, Catherine Lozupone, Sarah E Clark, William J Janssen, Christopher M Evans

## Abstract

T-helper (Th)-17 lymphocytes are central mediators of adaptive type 17 immunity. Decreased type-17 signaling increases severity of infections in humans and mice. However, detrimental effects of excessive type 17 responses in autoimmune and other inflammatory diseases highlight a need for type-17 immune calibration to support beneficial host defense requirements. Mechanisms of type 17 calibration are poorly understood. A gut-lung axis has been proposed to coordinate homeostatic protection and acute host defense. Factors that acutely alter the gut microbiome are heterogeneous and include acute intestinal infections, non-infectious colitis, and medical treatments such as antibiotics. How changes in the gut microbiome affect lung immune tone during homeostasis and acute pulmonary infections are also poorly understood. Prior studies have shown that antibiotics reduce expression of IL-17-mediated host defense in the gut. Since gut microbial homeostasis influences Th17 cell numbers in both the intestine and remote tissues, we postulated that antibiotic treatment would result in gut dysbiosis and weakened type-17 host defense in the lungs. We found that amoxicillin induces significant dysbiosis that is long-lasting and that there is a long-term decrease in type-17 tone in the lungs. We also found that in mice lacking the gut mucin, Muc2, Th17 cells increased in the lungs following inflammatory challenge. These findings suggest that antibiotic-induced dysbiosis can decrease lung immune defenses for long periods of time after cessation of antibiotic treatment.

## Introduction

Interactions between the microbiome and immune system in the gut are abundant, and studies demonstrate interdependence between gut microbial composition and gut Th17 signaling^1–6^. However, long-term effects of antibiotic induced gut microbiome changes on Th17 tone in the lungs have not been well characterized. T-helper (Th)-17 lymphocytes are central mediators of adaptive type-17 immunity, which is instrumental in mucosal defense against pathogens. Th17 cells are characterized by the expression of IL-17 isoforms including IL-17A and IL-22^7^. Type-17 signaling appears to be finely tuned as increased tone is implicated in the development of autoimmune disease^8^, whereas decreased tone has been associated with increased mortality from infections such as pneumonia^9,10^.

A decreased number of Th17 cells (but not Th1 cells) in bronchoalveolar lavage fluid from critically ill patients is associated with ventilator associated pneumonia^11^. Th17 cell numbers are decreased in the lungs of mice as a result of influenza infection, resulting in increased susceptibility to post-viral *Staphylococcus aureus* pneumonia^10^. In mice exposed to antibiotics, decreased IL-17A expression in the lungs is associated with increased mortality in *Streptococcus* pneumonia infections^9^. Antibiotic exposure is also associated with at least a 90-day period of heightened susceptibility to infections in humans, including pneumonia, and associated with increased mortality from bacterial pneumonia^9,12,13^. However, the mechanisms by which antibiotics affect levels of homeostatic type-17 signaling and the degree of acute response to subsequent insults in the lungs are poorly understood.

One way in which the gut microbiome can calibrate local Th17 cell numbers is via interactions between microbes and gut immune cells. A well characterized form of immune modulation is the interaction between segmented filamentous bacteria (SFB) – a gut commensal – and Th17 cells. SFB are gram positive, spore forming obligate anaerobes that reside in the terminal ileum and, unlike most other commensals, intimately associate with gut epithelial cells without causing overt inflammation^14^. SFB directly engage the Th17 T-cell receptor and induce the proliferation of Th17 cells^1,14^. Numerous studies have demonstrated that Th17 cell numbers in the gut are tightly correlated with the presence of SFB. In humans, SFB is most abundant during infancy and decreases in population size over time^15^. In mice, the presence of SFB is largely dependent on local microbiome compositions which can vary across breeding facilities and vendors^16^.

The relationship between the gut microbiome and the immune system can also manifest in other organs such as the lung, commonly termed the gut-lung axis^17^. While it is clear that gut microbes contribute to gut immune changes, less is known about gut microbial influences on lung immune signaling and mechanisms by which antibiotics disrupt these interactions. Lymphocytes, such as ILC2s^18^, Tfh17 cells^19^, and Th17 cells, can traffic from the gut to distant tissues, including the lung^1,20^, suggesting one possible mechanism for how the immune makeup of the gut can contribute to homeostatic immune tone in other tissues. However, there are currently no studies assessing whether changes in lymphocyte trafficking or lymphocyte numbers persist long-term after antibiotic cessation.

Our aim was to understand the long-term effects of antibiotic-induced gut dysbiosis on type-17 immunity in the lungs. Studies have shown that compared to other classes of antibiotics, low dose penicillin was more effective at decreasing SFB, IL-17A expression, and Th17 cell differentiation in the gut^2^. We exposed mice to a penicillin-type antibiotic, amoxicillin, and tested type-17 homeostatic and induced lung immune tone in the gut and lungs three weeks after cessation of antibiotic treatment. Our data show that amoxicillin treatment induces gut dysbiosis and a decrease in type-17 immune tone in the lungs, and that these effects are long-lasting. We also show that mice lacking a gut mucus barrier have increased numbers of Th17 cells in their lungs in response to sterile inflammation.

## Results

### Amoxicillin treatment alters the gut microbiome

We developed a mouse model to study the long-term effects of amoxicillin treatment on type-17 immune responses in the gut and the lungs. First, we tested how amoxicillin treatment affects the gut microbiome and whether the microbiome recovers 3 weeks after antibiotic treatment. Fecal samples were collected for microbiome evaluation prior to amoxicillin treatment (Day 0), on the last day of amoxicillin treatment (Day 21) and three weeks after cessation of amoxicillin (Day 42, Figure 1A). Control mice did not receive antibiotics, and their fecal samples were collected at the same time points.

**Figure 1:**
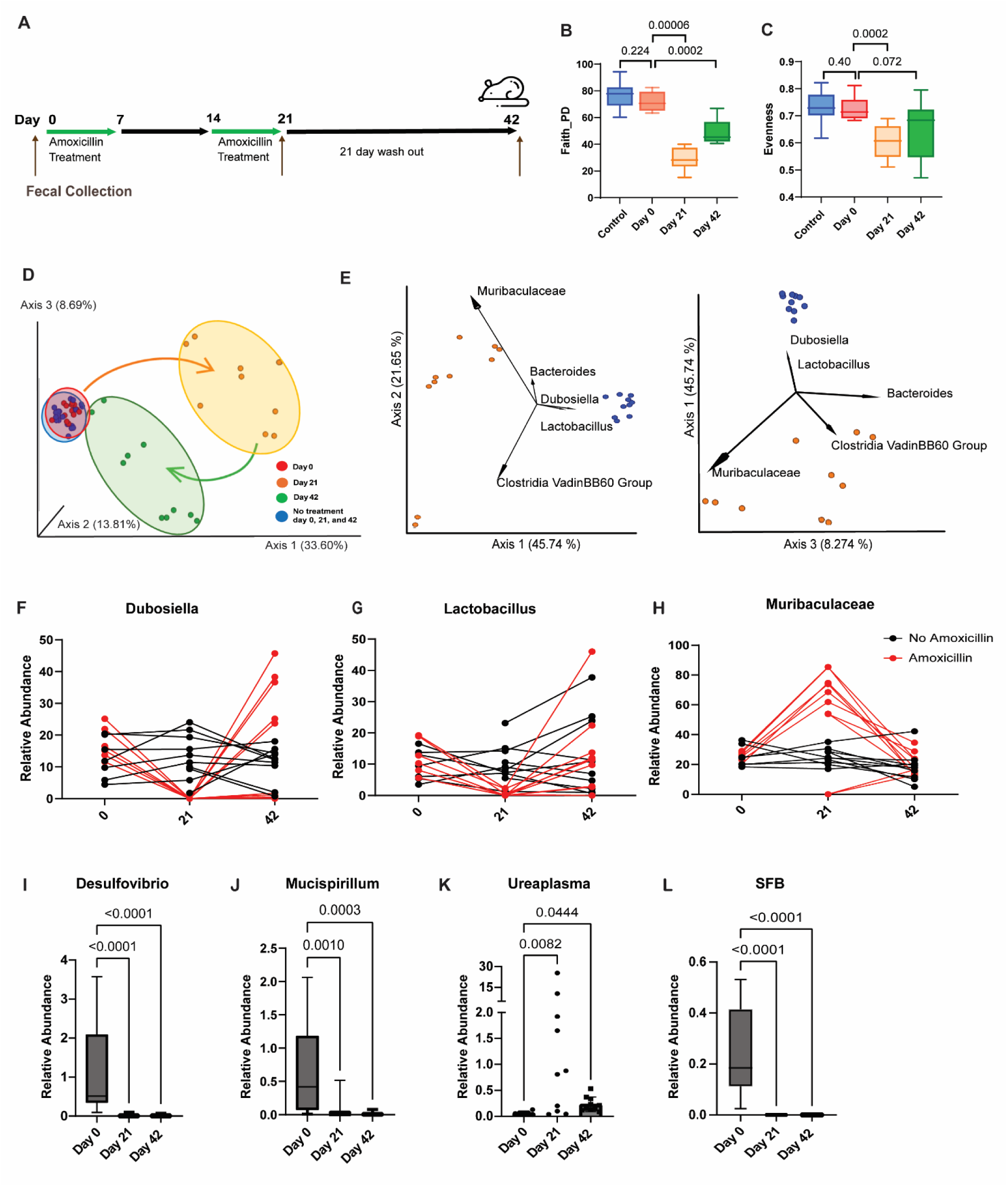
Amoxicillin induces gut dysbiosis: 8-week-old mice were treated with amoxicillin in their drinking water for 7 days, followed by a 7 day break and then an additional 7 days of amoxicillin. Fecal collection occurred on Day 0 – prior to amoxicillin initiation, Day 21 – final day of amoxicillin treatment, and day 42 – 3 weeks after cessation of antibiotics (A). 16S ribosomal DNA (rDNA) sequencing was used to assess microbiome changes, including alpha diversity by Faith’s PD (B) and Peilou’s evenness (C) as well as beta diversity by unweighted UniFrac (D) across all samples. PCoA biplots show the bacterial genera that are driving the clustering patterns on Day 21 (E, amoxicillin treated in orange and control in blue). Relative abundance for 3 of the dominant species identified in the PCoA plots over the 6-week time course (amoxicillin treated group in red and the control group in black) for *Dubosiella* (F), *Lactobacillus* (G), and *Muricaculaceae* (H). Relative abundance changes over the 6-week time course for the 3 bacterial genera identified in the ANCOM analysis as statistically different on Day 42 in the amoxicillin treated mice are plotted for *Desulfovibrio* (I), *Mucispirillum* (J), and *Ureaplasma* (K). Relative abundance for segmented filamentous bacteria (SFB) across the time course in amoxicillin treated animals (L). Significant differences in alpha diversity were determined using Kruskal-Wallis testing. To assess differences in relative abundance in the amoxicillin treated mice at multiple time points, Krustal-Wallis test with Dunn’s multiple comparisons test was applied, comparing Day 0 values to Day 21 and Day 0 to day 42. N=9-10 mice per group at every time point.

First, we assessed mouse gut microbiome community composition by measuring alpha and beta diversity. Mice treated with amoxicillin had significant reductions in the alpha diversity of their gut microbiomes at Day 21, as assessed by Faith’s phylogenetic diversity (Faith PD)^21,22^ and Peilou’s evenness^23^ (Figure 1B, C). Return to baseline alpha diversity remained incomplete at Day 42 by Faith’s PD (Figure 1B). Fecal samples from all time points from mice that were not exposed to antibiotic treatment were grouped together and showed no difference in alpha diversity compared to the Day 0 samples (Figure 1B, C, blue bars versus red bars).

To assess beta diversity, the unweighted UniFrac metric was used^24^. Principle coordinate analysis (PCoA) of pairwise UniFrac values showed separation between amoxicillin treated and untreated mice (Figure 1D, orange dots versus red dots) on Day 21, and these were significantly different with a PERMANOVA test with multiple test correction (False Discovery Rate (FDR) q=0.001). Three weeks after cessation of antibiotics, beta diversity partially returned to baseline, but there was still separation of the microbiome on Day 42 compared to Day 0 (Figure 1D, green dots compared to red dots) and a significant difference in overall community composition based on PERMANOVA of unweighted UniFrac distance values (FDR q=0.001). As with alpha diversity, fecal samples from all time points from mice not exposed to antibiotics were grouped together and compared to Day) samples, did not show significant differences in overall community composition based on PERMANOCA of unweighted UniFrac distance values (blue dots versus red dots, FDR q=0.3).

Next, we were interested in understanding which bacterial species were driving the differences in beta diversity with amoxicillin treatment (Day 21). We used the PCoA biplot functionality of QIIME 2^25^ to overlay bacterial genera in the same UniFrac PCoA space based on their relative abundances and plotted the 5 genera with the greatest distance from the origin. The genera *Dubosiella* and *Lactobacillus* were associated with controls and *Muricaculaceae* was associated with Amoxicillin treatment (Figure 1E). We then plotted the relative abundance over the time course in three of the species responsible for clustering at Day 21 and Day 42 that represented different patterns of recovery. While the relative abundances of these genera were stable over time in untreated mice, significant shifts were observed in antibiotic treated animals. For *Dubosiella* and *Lactobacillus* amoxicillin treatment caused significant decreases in relative abundance at Day 21 with rebound at Day 42 (Figure 1 F and G). For *Muribaculaceae*, there was an increase in relative abundance on Day 21, followed by decreasing levels on Day 42 (Figure 1H).

Microbiome sequencing data reports relative abundances, making it difficult to determine significance in abundance changes between samples. Analysis of Compositions of Microbiomes (ANCOM) is a statistical tool that allows for identification of differentially abundant taxa despite known microbiome sequencing constraints^26^. Using ANCOM, genus level microbiome differences were assessed on Days 0, 21 and 42 to further measure the effect of amoxicillin treatment on the change in relative abundance of specific genera. Prior to antibiotic treatment, there were no significant differences between the groups (data not shown). On the last day of amoxicillin treatment (Day 21), there were significantly more *Dubosiella* and *Lachnospiraceae NK4A136 group* species in mice that did not receive amoxicillin versus controls (Supplementary Figure 1A). Three weeks after cessation of amoxicillin, there were still significant differences detected by ANCOM between the amoxicillin treated and untreated groups. *Desulfovibrio* and *Mucispirillum* were significantly decreased in the mice that were treated with amoxicillin while *Ureaplasma* was elevated in the amoxicillin treated mice (Supplemental Figure 1B). To further evaluate the genus level shifts at Day 42 that were identified in the ANCOM analysis, relative abundance was plotted in the amoxicillin treated mice over the 6-week time course. *Desulfovibrio* and *Mucispirillum* levels dropped to undetectable levels and did not recover by Day 42 (Figure I and J). In comparison, *Ureaplasma* levels expanded on Day 21 and remained elevated on Day 42 compared to pre-treatment (Figure 1K).

Since SFB are known to induce Th17 cells in the gut, we evaluated amoxicillin effects on the relative abundance of *Candidatus arthromitus,* the predominant mouse strain of SFB. SFB were detectable at Day 0 but were no longer detectable on Day 21 after treatment with amoxicillin and remained undetectable on Day 42 after 3 weeks off antibiotics (Figure 1L).

### Amoxicillin treatment suppresses homeostatic type-17 immune tone in the terminal ileum and the lungs

We then assessed the long-term effects of amoxicillin induced gut dysbiosis on homeostatic type-17 gene expression in the gut and lungs. We treated mice with amoxicillin as described above. Twenty one days after antibiotic cessation (Day 42), type-17 cytokine gene expression was measured in the colon, terminal ileum and lungs of mice and compared to non-antibiotic treated littermate controls (Figure 2A). While there were no differences in gene expression of type-17 cytokines in the colon 21 days after antibiotics were discontinued (Figure 2B), there was a significant decrease in expression of *Il17a*, *Il22,* and *Ifnγ* in the terminal ileum (Figure 2C). In the lungs, expression of *Il17a* and *Il6*, both considered type-17 cytokines, was decreased 21 days after cessation of antibiotics whereas expression of *Ifnγ*, a type-1 cytokine, was unchanged (Figure 2D). Homeostatic IL-22 expression in the lungs was below the level of detection. Levels of *Il17a* gene expression in the lungs correlated with *Il17a* expression in the terminal ileum where SFB reside, but not in the colon (Figure 2E).

**Figure 2:**
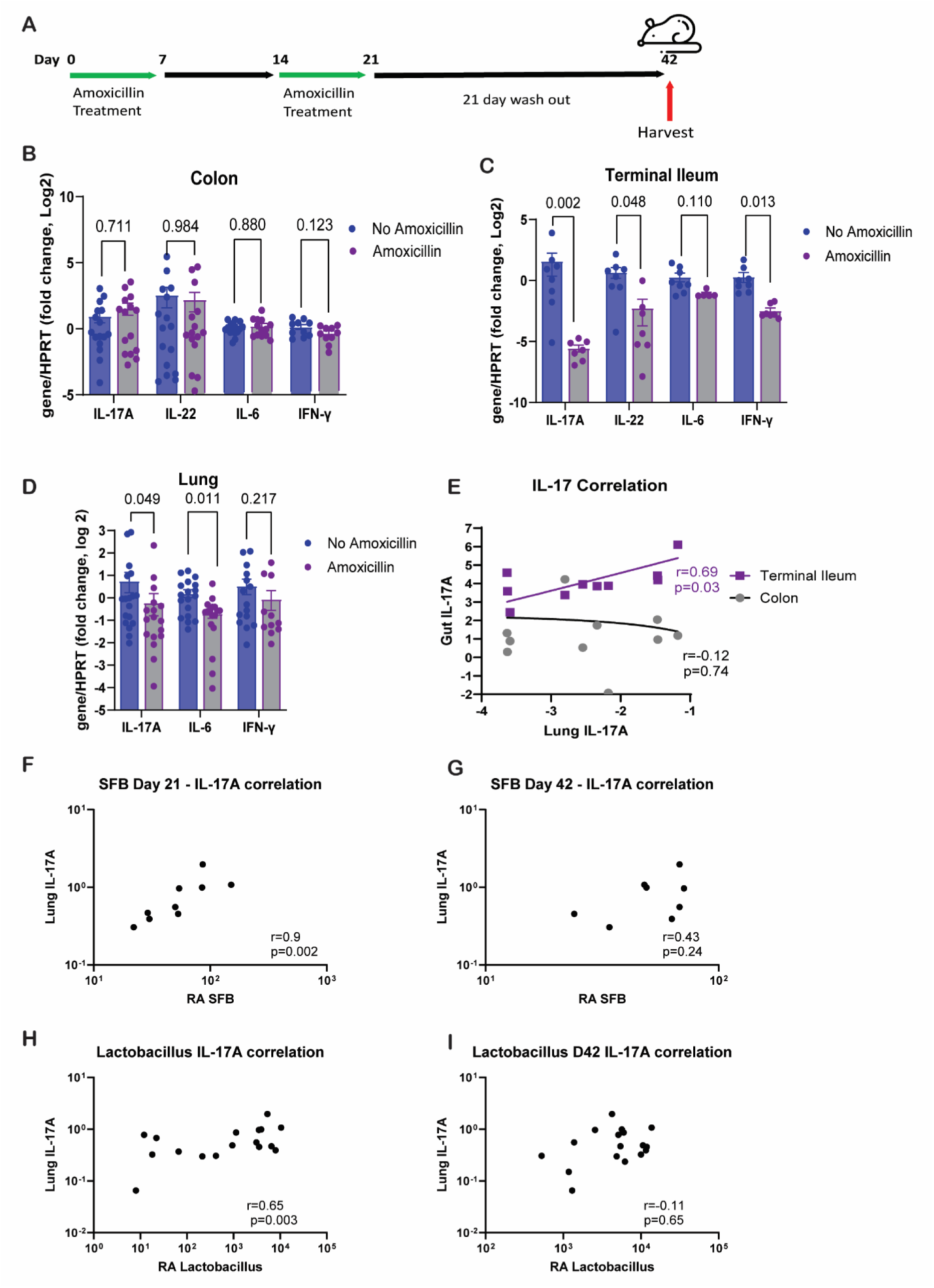
Amoxicillin induces long-term suppression of homeostatic type-17 lung immune tone: 8-week-old mice were treated with amoxicillin and tissues were harvested on day 42, which was 21 days after amoxicillin cessation (A). RNA was isolated from tissues and gene expression of IL-17A, IL-22, IL-6 and IFN-γ was evaluated by QPCR in the colon (B), terminal ileum (C), and lung (D) on Day 42. Blue circles/bars are the control animals and the purple circles/gray bars are the amoxicillin treated animals, N=7-16 per group, 2-4 experimental replicates performed. Data shown are mean ± SEM. Correlation of *Il17a* gene expression in the lungs was plotted against expression of *Il17a* in both the colon (gray line) and the terminal ileum (purple line) of the same animals on Day 42 (E). To determine correlation between bacterial species on different days and lung *Il17a* gene expression, lung *Il17a* expression on Day 42 was plotted against the relative abundance of SFB on Day 21 (F) and Day 42 (G). Similarly, lung *Il17a* expression on Day 42 was plotted against the relative abundance of Lactobacillus on Day 21 (H) and Day 42 (I). Statistical analysis was performed for every gene between groups using Mann-Whitney. Correlation significance was determined using Pearson coefficient.

Next, we sought to determine if the relative abundance of SFB in fecal material collected at different time points correlated with *Il17a* gene expression in the lungs. Because amoxicillin treatment reduced SFB levels below the level of detection, we focused on mice that did not receive antibiotics and had detectable SFB at all time points (blue dots from Figure 1B-E). Importantly, the relative abundance of SFB that was collected from non-antibiotic treated mice at the “Day 21” timepoint correlated very strongly with *Il17a* gene expression in the lungs twenty-one days later (spearman correlation coefficient (r)= 0.9, p=0.002, Figure 2F). However, the relative abundance of SFB in the fecal specimens collected on the “Day 42’ time point did not correlate with *Il17a* expression in the lungs on Day 42 (r=0.43, p=0.24, Figure 2G). To determine if bacteria other than SFB were potential contributors to changes in type-17 gene expression in the lungs, we tested if the 3 species of bacteria that helped define the PCoA clusters on Day 21 (last day of amoxicillin treatment) correlated with *Il17a* gene expression in the lungs on Day 42. As with SFB, there was significant correlation seen between *Il17a* gene expression in the lungs at Day 42 and the relative abundance of *Lactobacillus* species on Day 21 (Figure 2H), but not Day 42 (Figure 2I). *Dubosiella* and *Muricaculaceae* relative abundance at either time point did not correlate with *Il17a* gene expression (data not shown).

### Amoxicillin decreases Th17 cell numbers in the lungs during sterile inflammation

Next we assessed the effects of amoxicillin treatment on Th17 cell counts in the lungs. In the absence of inflammation, there were small numbers of Th17 cells in the lungs (<2% of total CD4 cells). Non-typeable *Haemophilus influenzae* lysate (NTHi) has previously been shown to induce lymphocyte recruitment to the lungs^27^ and increase Th17 cell signaling^28^. Thus, we challenged mice with aerosolized NTHi to induce sterile inflammation. WT mice were treated with amoxicillin as described above. On Day 42, amoxicillin and non-amoxicillin pre-treated mice were challenged with aerosolized NTHi and endpoints were collected 48 hours later (Figure 3A). Th17 cells were identified by flow cytometry from whole lung digests. In these experiments, CD4+ expressing T cells (CD45+CD3+CD4+) were classified as Th17 cells if they also expressed CCR6 and intracellular RORγT. Th1 cells were identified by expression of intracellular TBET and absence of RORγT (Figure 2B, gating strategy – Supplementary Figure 2). There was an increase in Th17 cell numbers in the lungs after NTHi challenge, but this increase was significantly blunted in animals that had been pre-treated with amoxicillin (Figure 3C). Amoxicillin pre-treatment did not change the frequency of Th1 cells or neutrophils at baseline or following challenge with NTHi (Figure 3 D,E). There was also no change in the number of γδ-T cells (another major source of IL-17A in the lungs) in any of the conditions (Supplementary Figure 3).

**Figure 3:**
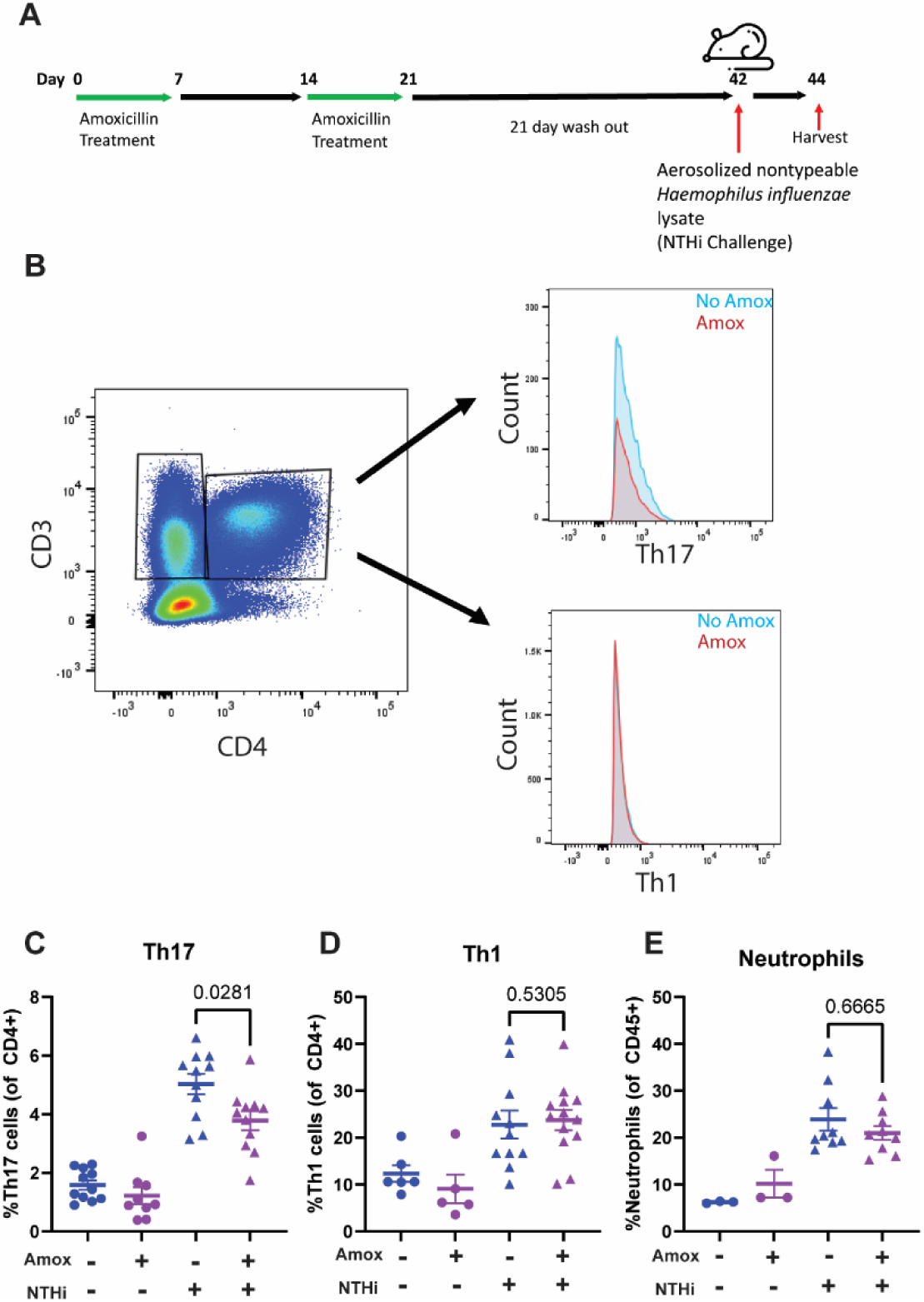
Amoxicillin induces long-term suppression of Th17 cell numbers in the lungs in response to sterile inflammation: 8-week-old mice were treated with amoxicillin and on day 42 were challenged with aerosolized, heat killed non-typeable *Haemophilus influenzae* lysate. Tissues were harvested 48 hours later (A). Lung tissue was harvested, digested and used for flow cytometry. CD45+CD3+CD4+ cells were further gated on Th17 cells (CCR6+ and RORγT+) or Th1 cells (TBET+), depicted as histograms. Control mice that did not receive amoxicillin are shown in blue histograms and amoxicillin treated mice are shown in red histograms (B). Flow cytometry was used to assess percentages of Th17 cells (C), Th1 cells (D), and neutrophils (E) in the lungs of mice that did not receive antibiotics (blue circles/triangles) compared to those that did (purple circles/triangles) both in the absence of NTHi (circles) and in the presence of NTHi (triangles). Significance between amoxicillin treated and untreated mice in the NTHi exposed group was determined using a Mann-Whitney test. N=9-11, 3-4 replicates performed per condition.

### Lung Th17 cell numbers increased in Muc2 KO mice

Next, as a gain-of-function experiment, we used a mouse strain that lacks the *Muc2* gene as these mice are prone to increased Th17 tone in the gut^29,30^. Muc2 absence results in loss of intestinal mucus barriers, which in turn predisposes mice to colitis^31^. We tested if the increase in type-17 tone in the gut resulted in increased type-17 tone in the lungs during sterile inflammation. We exposed *Muc2^-/-^* mice and WT littermates (age-matched to the amoxicillin treated mice) to aerosolized, heat-killed NTHi and then 48 hours later measured Th17 cell numbers in the lungs by flow cytometry. There was a significant increase in Th17 cells in the lungs of the *Muc2^-/-^* mice 48 hours after NTHi exposure (Figure 4A). However, there was no difference in Th1 cells or neutrophils (Figure 4 B,C). We did not see differences in γδ-T cell, NK or NK T-cell populations between the *Muc2^-/-^* and *Muc2^+/+^* mice at baseline or after NTHi challenge (Supplementary Figure 4).

**Figure 4:**
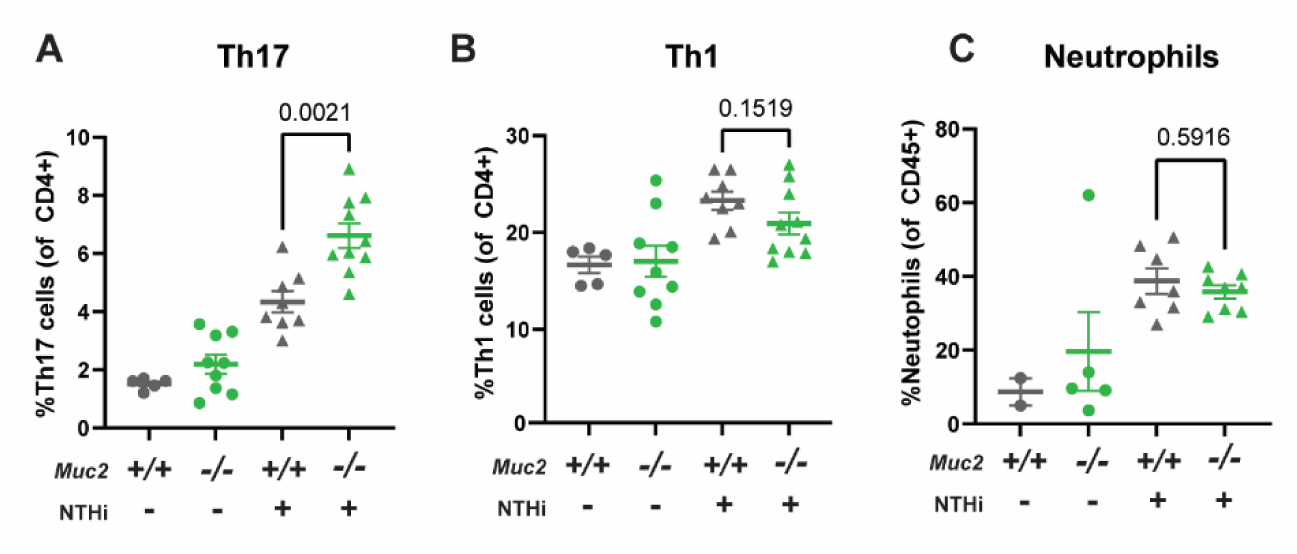
*Muc2^-/-^* (KO) mice have increased Th17 lung immune tone: At 12-16 weeks of age, Muc2 WT and KO mice were exposed to NTHi and tissues were harvested 48 hours later. Lung digests were used for flow cytometry to assess percentages of Th17 cell (A), Th1 cell (B), and neutrophil (C) numbers. Significance between *Muc2^+/+^*and *Muc2^-/-^* mice in the NTHi exposed group was determined using a Mann-Whitney test. N=5-10, 2 replicates per condition were performed.

### Fecal reconstitution after amoxicillin treatment restores type-17 signaling in the gut and lungs

To test if the changes in type-17 lung tone could be rescued by restoring gut microbiome composition, we performed fecal reconstitution experiments. 24 hours after mice completed amoxicillin treatment they received fecal transplant with fecal material from healthy, age-matched non-antibiotic exposed donor mice for 3 days in a row. Control mice received amoxicillin but no microbiota transfer. Mice were challenged with NTHi 21 days later (Figure 5A). First, we compared microbiome recovery between the mice that received fecal transplants (on days 22-24) to untransplanted controls. Microbiome analysis showed that alpha diversity in mice that received fecal transplant after antibiotic treatment was restored by Day 42 (Figure 5B) compared to mice that received amoxicillin treatment alone. Similarly, beta diversity was restored by fecal transplantation by Day 42 (Figure 5C, light green dots) as opposed to the mice that received amoxicillin alone. Next, we assessed the relative abundance of SFB with and without fecal reconstitution. While the mice that did not receive amoxicillin showed stable levels of SFB over the 6-week experiment, the mice treated with amoxicillin showed reduced levels of SFB at Day 21 with no recovery by Day 42 (Figure 1M, 5D). The mice that received fecal transplant after amoxicillin treatment showed restored levels of SFB at Day 42 (Figure 5D). Bacterial genus level analysis showed similar bacterial families, such as *Dubosiella* and *Lachnospiraceae NK4A136 group,* were responsible for the clustering seen in beta diversity as before (Supplementary Figure 5).

**Figure 5:**
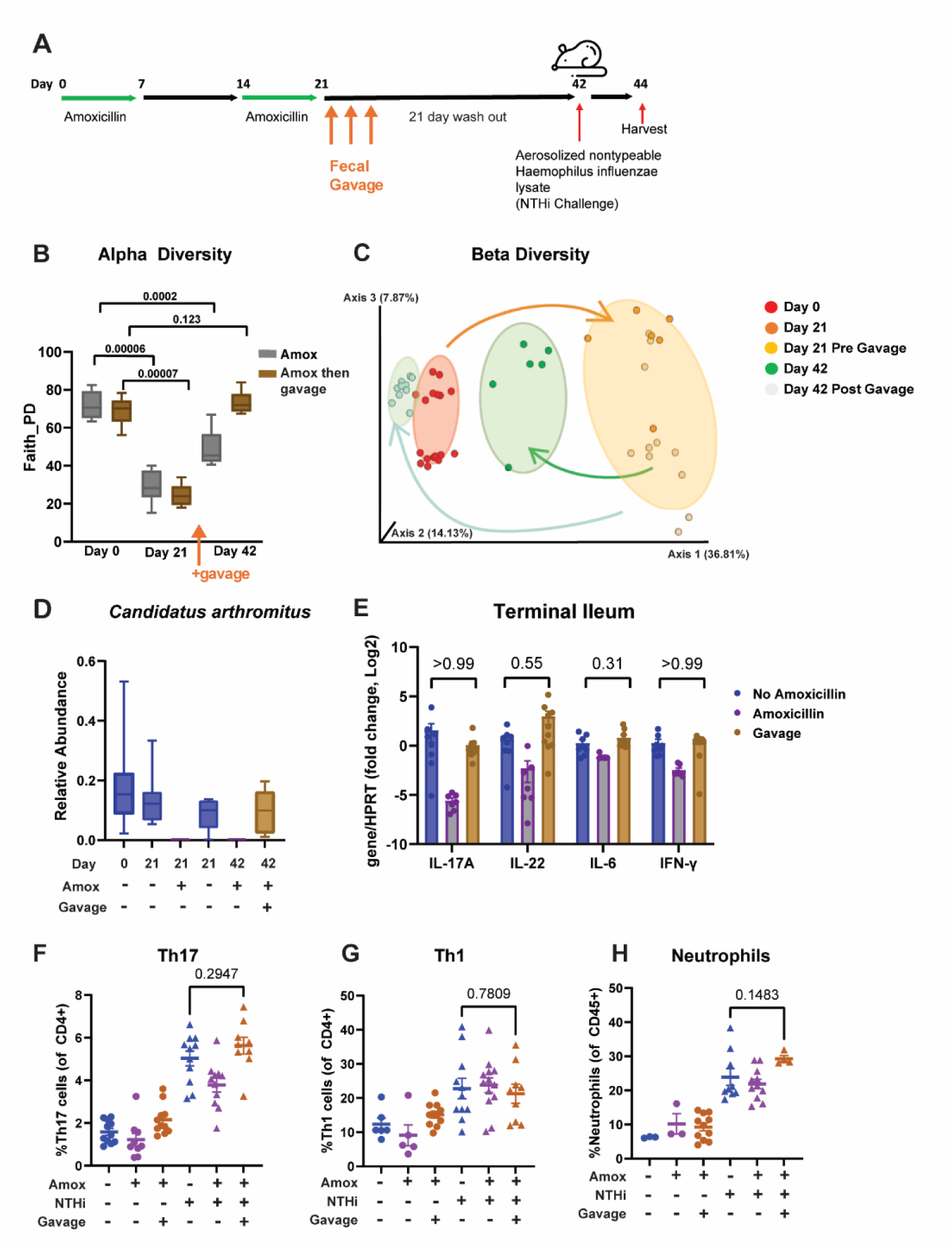
Fecal Reconstitution After Amoxicillin Treatment Restored Type-17 Immune Tone: 8-week old mice were treated with amoxicillin as before. Fecal material was collected from mice not receiving antibiotics and used to gavage the amoxicillin treated mice daily starting on day 22 for a total of 3 days. On day 42, tissues were harvested or mice were exposed to aerosolized NTHi (A). Fecal pellets were collected on days 0, 21, and 42. 16S ribosomal DNA (rDNA) sequencing was used to assess microbiome changes over time, including alpha (B) and beta (C) diversity in the mice that received amoxicillin only (orange dots and dark green dots) compared to mice that received amoxicillin followed by fecal gavage (yellow dots and light green dots). Relative abundance of segmented filamentous bacteria was assessed in all groups (control, amoxicillin treated, and amoxicillin treated then gavaged) at all time points (Day 0, Day 21, and Day 42) (D). Terminal ileum was harvested on day 42, RNA was isolated and QPCR was performed to quantify expression of *Il17a*, *Il22*, *Il6* and *Ifnγ*. Blue bars are the control, no amoxicillin group, purple bars are the amoxicillin treated group, and the brown bars are the amoxicillin treatment followed by fecal gavage group (E). Lungs were harvested either on day 42 for baseline measurements or 48 hours after NTHi challenge for flow cytometry analysis of Th17 cells (F), Th1 cells (G), and neutrophils (H). Non-antibiotic treated mice are in blue, amoxicillin treated mice are in pink and amoxicillin treated followed by fecal gavage mice are in brown. Circles are in the absence of NTHi and triangles are in the presence of NTHi. Statistical significance between the antibiotic untreated mice and the mice that were treated with amoxicillin followed by fecal gavage was assessed using a Mann-Whitney test. N=7-17, at least 2 replicates were performed for each condition.

Next, we assessed type-17 tone in the terminal ileum and lungs with and without fecal gavage after amoxicillin treatment. Gene expression of type-17 cytokines (IL-17A and IL-22) was restored in the terminal ileum on Day 42 in the mice receiving fecal gavage (Figure 5 E). We then assessed Th17 cell numbers in the lungs after sterile inflammation. Mice were exposed to inhaled NTHi as before, and Th17 cell percentages were measured in the lungs 48 hours later by flow cytometry. Th17 cell percentages were restored in mice that received fecal gavage (Figure 5F). Th1 and neutrophil cell numbers were not affected by amoxicillin treatment or fecal gavage (Figure 5 G, H). Taken as a whole, these data show that gut microbiome reconstitution was able to restore type-17 tone in both the gut and the lungs.

## Discussion

Our data show that amoxicillin is associated with gut dysbiosis in mice leading to long-lasting suppression of type-17 immune tone in the small intestines and in the lungs. We also show that depletion of mucus in the gut results in increased type-17 immune tone in the lungs during inflammation. This is one of the first studies to show the long-lasting effects of antibiotic treatment on immune populations in the gut and the lungs as well as one of the first to demonstrate a change in type-17 immune phenotype in the lungs of *Muc2^-/-^* mice in the setting of sterile inflammation.

Many studies exploring the effects of antibiotics on gut and lung immune responses use a 4-drug antibiotic cocktail, such as ampicillin, neomycin, metronidazole, and vancomycin^32^. It is usually administered for 10-14 days, and end points are collected 1-3 days after cessation of antibiotics. In our studies, we used amoxicillin monotherapy, we treated mice with a “pulsed” course, and we waited 21 days after antibiotic cessation prior to assessing immune responses.

In the outpatient setting, most people receive prescriptions for a single antibiotic for 5-10 days duration. Amoxicillin is one of the most prescribed antibiotics in the United States and was therefore the agent we chose for our study^33^. Importantly, up to 20% of patients receive a second prescription for antibiotics within 10 days for the same outpatient illness^34^. Thus, treating mice with repeat amoxicillin monotherapy is more clinically relevant than a 4-drug regimen. Studies have shown that there is a tendency for fungal overgrowth if antibiotics are administered for more than 7 days^35^. To avoid fungal overgrowth, we opted for a shorter antibiotic treatment duration of 7 days as opposed to the commonly used 10-14 days. Also, antibiotics administered as pulsed doses (one week of antibiotics followed by a one-week break and then an additional week of antibiotics) were more effective than continuously dosed antibiotics at decreasing Th17 cell populations in the gut^36^. For all these reasons, we treated our mice with amoxicillin for 7 days, followed by a 7-day break, and then an additional 7 days of amoxicillin.

In this study, we chose the 3-week time point because this is when trained innate immune responses and adaptive memory responses are established in mice^37,38^. As such, we consider this a “long-term” timepoint. The studies assessing effects of antibiotic pre-treatment on bacterial pneumonia outcomes in mice generally waited 1-3 days prior to bacterial challenge to allow time for the antibiotics to wash out so as not to affect outcomes^9^. As patients are at increased risk of infections for up to 90 days following antibiotic treatment^13^, we were interested in understanding the long-term effects of antibiotic treatment on both microbiome recovery and lung immune responses; thus, we waited 21 days after antibiotic cessation to collect our end points.

An important observation in this study is that the microbiome did not completely recover 3 weeks after cessation of antibiotics. Some bacterial species, such as *Desulfovibrio, Mucispirillum,* and SFB did not recover at all three weeks after antibiotic cessation, whereas *Ureaplasma* recovered to higher relative abundance compared to the pre-antibiotic time point. Other species, such as *Lactobacillus, and Muribaculaceae* recovered to baseline 3 weeks later. These findings support a recently published hypothesis that the human gut microbiome reverts to a new baseline after antibiotic treatment^39^. As in our study, a human study assessing longer-term recovery after antibiotics showed that some low abundance bacterial species do not recover, even after 180 days^40^. Even low abundance taxa, such as SFB, can contribute significantly to immune signaling in multiple tissues, including the gut and the lungs. While antibiotic treatment is a well-recognized risk of *Clostridioides difficile* infection, less is known about long-term risks associated with sustained microbiome changes. Collectively, these studies contribute to the rising concern that even short courses of antibiotics can have long-lasting effects on bacterial microbiome compositions with potentially deleterious consequences to homeostatic immune responses in distant organs.

While our study targeted SFB with amoxicillin treatment, it is clear that amoxicillin has other effects on the gut microbiome. Some of the bacterial communities that were most affected by amoxicillin treatment have been previously associated with chronic lung and autoimmune diseases^41–43^, suggesting their clinical relevance in contributing to immune tone at distant sites. This study highlights that antibiotic-induced dysbiosis is long-lasting and could contribute to other disease states at distant time points. Further studies are necessary to understand how long-lasting these effects are on our microbiomes and if homeostatic and induced lung immune responses are impaired long term.

The observation that SFB numbers on Day 21, but not Day 42, correlated significantly with *Il17a* gene expression in the lungs on Day 42 (spearman correlation coefficient 0.9, p=0.002, Figure 2F) has important possible implications. While we did not assess gene expression in the lungs in our study on Day 21, it is surprising that relative abundance of SFB on the same day as lung harvest had no correlation to *Il17a* gene expression. We saw a similar pattern for *Lactobacillus* – relative abundance on Day 21, but not Day 42, correlated to the levels of *Il17a* gene expression in the lungs on Day 42. This suggests that a snapshot of microbial composition on any given day does not necessarily represent the bigger picture of dynamic microbial shifts that contribute to distant organ immune phenotypes. How microbiome signals transmit to distant organs is still an active area of investigation. Our data suggests that some of the signals might not be transmitted instantly and could take days or weeks to transmit. It is currently unclear if this is because it takes time for cells to traffic or if the microbiome signals are affecting trained immunity, which takes time to develop. Further research is needed to understand the mechanism and timing of how gut microbial composition transmits signals to distal tissues.

Dysbiosis has also been associated with inflammatory bowel disease (IBD). A large percentage of patients with IBD also have pulmonary disease^44^, providing further support for the gut-lung axis. However, the mechanistic drivers of lung pathology in these individuals are poorly understood. In mice, a disruption of the gut mucus barrier can predispose to bowel inflammation and colitis^31^. MUC2 and MUC5B are the predominant mucins that support mucus defense in the gut and lungs, respectively. Mice lacking the mucin proteins Muc2 or Muc5b have disrupted mucus barriers and display dysbiosis in the gut and lung^45,46^. There is evidence that *Muc2^-/-^*mice have increased Th17 tone in the gut^29,30^. However, there are no studies investigating Th17 cell numbers in the lungs of *Muc2* deficient mice. Our data suggest that there is also an increase in Th17 cell numbers in the lungs of *Muc2* deficient mice when there is inflammation present.

One bacteria type that increases in relative abundance with amoxicillin treatment following antibiotic treatment is *Muribaculaceae.* Bacteria in this family can digest MUC2 as a source of energy^47^ and fecal microbiome studies in *Muc2^-/-^*deficient mice show gut dysbiosis with a decrease in the relative abundance of *Muribaculaceae*^48^. The relative abundance of this taxon increased on the last day of antibiotic treatment (Day 21, Figure 1I) in our study. If the relative abundance increase represents a bloom of these bacteria, then increased mucus degradation could also negatively impact the gut environment resulting in increased inflammation for the duration of antibiotic administration.

While our studies identify a significant role for Th17 cells in the gut-lung axis, precisely how changes in gut microbial composition are transmitted to the lungs remain poorly understood. The most likely mechanisms of gut-lung axis communication include trafficking of immune cells from the gut to the lung and/or changes in gut metabolites affecting distant organ systems, including the lungs. While there is compelling evidence that some cell types traffic from the gut to the lungs, evidence also suggests that gut microbial-derived metabolites circulate to distant organs^49^. As an example, short chain fatty acids (SCFAs) derived from bacteria enable crosstalk between the gut and other distant organs, such as the brain^50^. Investigating which gut microbiome-dependent metabolites enter the circulation and affect immune signaling at distant sites is an important next step in understanding how the gut-lung axis can be manipulated to improve barrier function and immune responses in the setting of infection.

In conclusion, our study shows that amoxicillin use results in long-term dysbiosis, decreased type-17 lung immune tone, and altered gene expression in the terminal ileum. We also show a lung phenotype in *Muc2* deficient mice with increased accumulation of Th17 cells in response to sterile inflammation in the lungs.

## Methods

### Mice

All studies were conducted with the approval of the Institutional Animal Care and Use Committees at the University of Colorado. Housing rooms were maintained at 22°C, 30-40% humidity, and a 14/10 (h/h) light dark cycle. All strains are on C57BL/6 backgrounds. Equal numbers of males and females were used.

*Muc5b*^mEGFP^ mice were generated by insertion of a monomeric enhanced green fluorescent protein (mEGFP) expression cassette into the third intron of the mouse *Muc5b* gene. The mEGFP sequence was flanked by splice acceptor (SA) and splice donor (SD) sequences that enable its introduction into *Muc5b* mRNA.

To introduce the SA-mEGFP-SD sequence, a double-strand in *Muc5b* intron 3 was introduced using CRISPR-Cas9 directed at the forward strand sequence 5’-GAACATCTGTTGACCACCCT. Homology directed repair (HDR) was mediated by additional 5’ and 3’ homology sequences in a 1,622 nt single stranded deoxyoligonucleotide (oligo). Homologous insertion was determined using PCR with primers targeting regions in mEGFP and external to the HDR oligo (“Left” side: Intron2 fwd: 5’-ACCCTTAGCTACATCCCCGTG, mEGFP rev: 5’- GCAGATGAACTTCAGGGTCAG; “Right” side: mEGFP fwd 5’-ATCACTCTCGGCATGGACGAG, Intron 4 rev 5’- AGAAAGCTCCGGTCTTCCTTC (Supplementary Figure 6A). Results were verified by sequencing of PCR amplicons.

Introduction of EGFP at this region of the Muc5b protein was reported previously^51^. Consistent with our prior result, we did not observe changes in Muc5b protein assembly or localization in the lungs (data not shown). *Muc5b*^mEGFP^ mice were produced on a C57BL/6 background and used as wild type for studies reported here.

*Muc2* knockout mice were generated using CRISPR-Cas9 to introduce DNA breaks in the promoter and intron 2 of the *Muc2* gene. CRISPR guides were directed to target sites using 5’-ACCTCACTTCACGTCATCCA and 5’-CCTATCCCACATGCCCCCCA sequences found on the reverse strand of *Muc2*. DNA repair mediated by non-homologous end-joining resulted in mutant alleles lacking core promoter and 5’ end coding sequences required for *Muc2* gene expression. This was confirmed by PCR (WT fwd: 5’-GATACCATCTCTTCTGCTCTG, WT rev: 5’- GCTTTCCTGGAACTGCCACAC; KO fwd: 5’-GACACGAGTCCAATGAGAGTTG, KO rev: 5’-GCTTTCCTGGAACTGCCACAC and DNA sequencing (see Supplementary Figure 6B).

Consistent with a prior line of *Muc2* deficient mice^52^, the line produced here demonstrated severe attenuation of goblet cell presence in the small and large intestines (data not shown). Also consistent with prior reports^52,53^, *Muc2*^−/−^ mice displayed abnormal bowel functions and early mortality. Reduced survival was apparent only after 20 weeks age (data not shown), so studies reported here employed younger mice (ages 12-16 weeks). *Muc2*^−/−^ were produced on a C57BL/6 background, bred by crossing heterozygotes, and used in experiments alongside wild type littermate controls.

### Amoxicillin Exposure

Starting at 8 weeks of age, mice were given amoxicillin (0.26 mg/ml) *ad libitum* in drinking water for 7 days, followed by a 7-day washout, and then 7 additional days of treatment.

### Microbiome transfers

2 g of fecal pellets from 14 non-antibiotic treated mice (from different cages and equal numbers of males and females) were suspended in 20 ml of PBS, agitated into a slurry, and frozen down at -20°C in 1 ml syringes that were capped to prevent exposure to air. On day of microbiome transfer, 1 ml of fecal slurry was thawed and mixed 1:2 with sterile PBS, filtered through 100 µm filter, and 200 ul of the diluted fecal suspension was gavaged into the antibiotic treated mice for 3 consecutive days starting 24 hours after antibiotics were discontinued.

### NTHi Exposure

5 ml of head-killed non-typeable *H. influenza* lysate (NTHi, protein concentration 2.5 mg/ml) was placed in an Ultravent nebulizer (Covidien) driven by 200 kPa in air for 30-40 minutes until all the lysate was aerosolized.

### Microbiome Data Generation

Briefly, total genomic DNA was extracted using the DNeasy PowerSoil Kit and the V4 variable region of the 16S rDNA gene was PCR amplified using the 515F:806R primers and standard protocols of the Earth Microbiome Project^54,55^. Amplified DNA was quantified in a PicoGreen (ThermoFisher Scientific) assay and equal quantities of DNA from each sample was pooled. The pooled DNA was sequenced using a V2 2x250 kit on the Illumina MiSeq platform (San Diego, CA) at the Anschutz Center for Microbiome Excellence. Sequencing data was analyzed using the QIIME2 (version 2023.5) bioinformatics platform. Statistical analysis described below.

### Bronchoalveolar Lavage and Lung Harvest

After euthanasia, the trachea was exposed, and an 18- gauge Luer stub was inserted into an incision made on the ventral portion and secured with silk thread. The lungs were lavaged by instilling and removing 1 ml PBS containing 0.6 mM EDTA three times (PBS/EDTA, 1 ml total). BAL fluid was used for hemocytometer counts and differentials. Following lavage, the lungs were perfused with 20 ml PBS via cardiac puncture, the right middle lobe was tied off and 1.25 ml of dispase (StemCell Technologies, 5u/mL) was instilled into the lungs. The trachea was tied off and the lungs were isolated. The RML was removed and placed into 0.5 ml RLT buffer and flash frozen on dry ice while the rest of the lungs were placed into 1 ml of dispase and incubated for 45 min at room temperature. The lungs were then minced into fine pieces with tweezers and passaged through a 70 µm filter, washed with media and passaged through a 40 µm filter. The cells were centrifuged for 10 min at 2,000 rpm at 15 °C, washed with cold MACS buffer and centrifuged again. The cells were incubated with EpCAM beads (Milteny) at 4°C for 10 minutes, washed with cold MACS buffer, and centrifuged again. The cells were resuspended in 1 ml cold MACS buffer and run through MACS columns per manufacturer’s instructions (Milteny). The flow through was used for flow cytometry (as below).

### Gut tissue harvest

After lungs were harvested, the abdomen was exposed and the colon was isolated. The fecal material was removed and 2.5 cm of the distal colon tissue was placed in RLT buffer and flash frozen on dry ice. Next, the terminal ileum was isolated and 2.5 cm of the terminal ileum just proximal to the cecum was collected and placed into RLT buffer and flash frozen on dry ice.

### QRT-PCR

Tissues harvested were placed in RLT buffer (Qiagen) and frozen at -80°C. Tissues were thawed, homogenized using an MPBio Fast Prep 24 grinder and lysis system, and RNA was isolated using Qiagen mini RNA kits. RNA quantity was determined by NanoDrop (ThermoFisher Scientific, Waltham, MA). 800 ng of RNA was used to make cDNA using the Thermo Scientific, SuperScript III first strand synthesis kit. QPCR was run on 2 µl of cDNA using Taqman Master Mix (Thermo FisherTaqMan Fast Advanced Master Mix) and the following TaqMan probes IL-6 (Mm00446190_m1), IL-17A (Mm00439618_m1), IL-22 (Mm01226722_g1), IFN-γ (Mm01168134_m1), and HPRT (Mm03024075_m1). QPCR was run on QuantStudio 7 Flex (Applied Biosystems).

### Flow Cytometry

EpCAM- cells were centrifuged at 300g for 5 min, washed in PBS and stained with either the lymphocyte panel or the monocyte panel. For the lymphocyte panel, the cells were stained with live/dead stain for 30 min at room temperature in the dark, then washed with FACS buffer, centrifuged at 300g for 5 min, and stained for the following surface markers in the presence of Fc Block (Clone 93, Biolegend) at a dilution of 1:100: CD45, CD3, CD8, CD4, CCR6, TCRγδ, and DX5 for 30 min at room temperature. The cells were then washed in FACS buffer, centrifuged at 300g for 5 min, resuspended in 1 ml of Fix/perm buffer (ThermoFisher, catalogue #00-5523-00), and placed at 4°C overnight. The next day, the cells were washed in perm buffer wash, centrifuged at 300g for 5 min, and stained for intracellular markers at a dilution of 1:100: RORγT and TBET for 30 min at room temperature. The cells were washed in perm wash buffer, centrifuged at 300g for 5 min, resuspended in 1 ml of FACS buffer and acquired on the BD LSR Fortessa analyzer. For the monocyte panel, the cells were incubated with Fc block for 15 min on ice and then stained with the following surface markers at a dilution of 1:100: CD3, CD45, CD11b, CD11c, Siglec-F, Ly6g, Ly6C, and F4/80 for 30 min at room temperature. The cells were washed with FACS buffer, centrifuged at 300g for 5 min, resuspended in 1 ml FACS buffer and stained with 1 µl DAPI (1 mg/ml) prior to acquisition on the BD LSR Fortessa analyzer

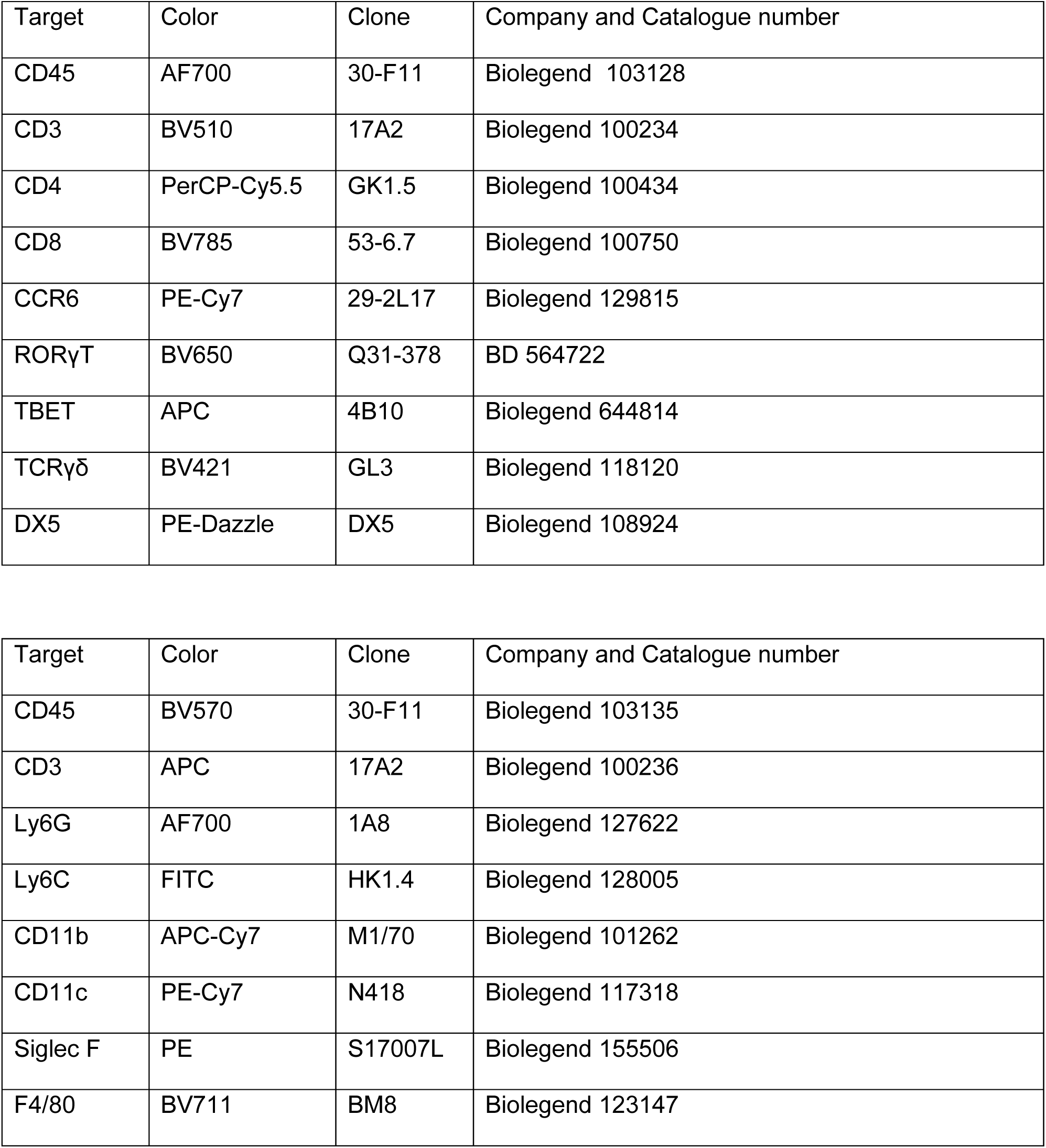

### Statistical Analysis

#### Microbiome

The following 16S data processing and microbiome analysis were performed using QIIME2 (version 2024.2)^25^. FASTQ files were demultiplexed and then denoised and identical sequences binned into identical Amplicon Sequence Variants (ASVs), using DADA2^56^. ASVs were assigned taxonomically using the Naïve Bayes classifier trained on SILVA 138(silva-138-99-515-806-nb-classifier and _sepp-refs-silva-128). A phylogenetic tree was built using the SEPP plugin^57^. Features that did not classify at the phylum level or were classified as mitochondria or chloroplast were filtered from the analysis. Samples were rarefied at a depth of 23,328. Alpha diversity was evaluated by using Faith’s Phylogenetic Diversity^22^ and Pielou’s Evenness^23^. Kruskal-Wallace analysis was used to determine statistical significance. Beta diversity was assessed using unweighted UniFrac distance^24^. Beta diversity was assessed using weighted and unweighted UniFrac distance and these distances were used to generate Principal coordinate analyses (PCoA) and biplots at taxonomic level 6 (genus). To do this importance of each genus was calculated for each feature based on the vector’s magnitude (euclidean distance from origin) and the genera with the 5 largest importance scores were plotted in the same UniFrac PCoA space using Emperor^58^. Permutation-based multivariate analysis of variance (PERMANOVA) was performed to test for associations between microbial community composition and variables of interests and potential cofounders including Sex, Day, Cage ID, and Treatment Day. Analysis of Composition of Microbiomes (ANCOM) was performed on data stratified by timepoint (Day 0, Day 21, Day 42) on taxonomic level 6. Unweighted biplots were also created using data stratified by timepoint at taxonomic level 6 to visualize the top taxa, as identified by biplot, as vectors within the PCoAs.

#### Gene expression

As this was not normally distributed data, Mann-Whitney tests were used for two sample comparisons, and Kruskal-Wallis tests were used for more than two sample comparisons.

#### Immune populations

As this was not normally distributed data, Mann-Whitney tests were used for two sample comparisons, and Kruskal-Wallis tests were used for more than two sample comparisons.

## Acknowledgements

We would like to thank the Genetically Engineered Murine Models (GEMM) Core at the University of Colorado (RRID:SCR_025491) for assistance in creating the Muc5b-GFP and Muc2 knockout mouse lines. The authors would like to thank the NIH for the following funding sources: R01HL080396, R01HL179623, I01BX005343, P01HL162607 (CME); R01HL14974, R35HL140039 (WJJ); R01HL130938 (CME, WJJ); 5T32HL007085, 1F32HL172725, P30 AR079369 (MO), DK131581 (CL, BP), R03 DK114545 (CMA).

## Supplementary Figures

**Supplementary Figure 1:**
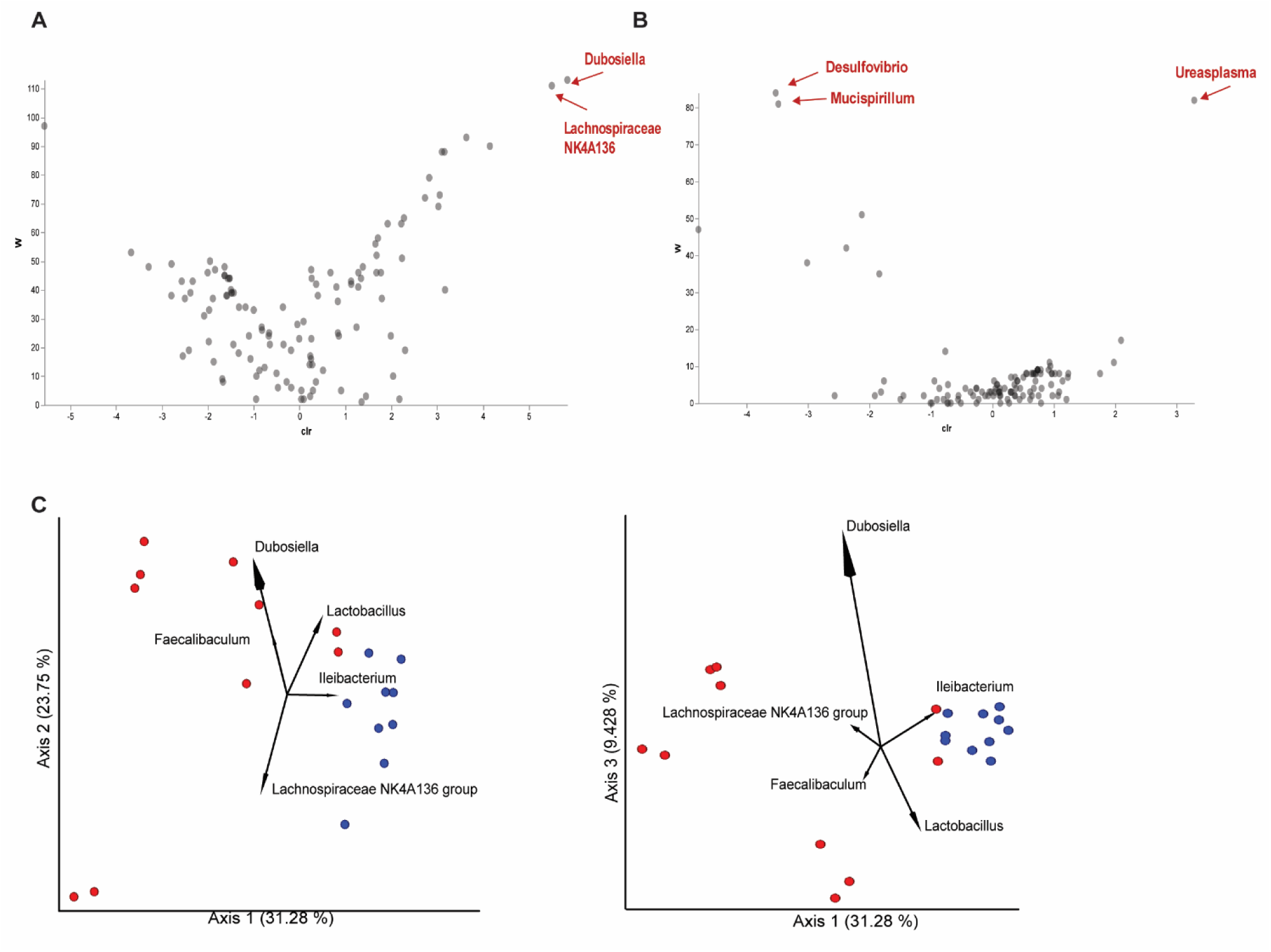
Amoxicillin induces gut dysbiosis is not fully resolved by Day 42: ANCOM analysis of microbial diversity in the amoxicillin treated and untreated groups. Volcano plot showing differential abundance of bacteria on Day 21 between the two groups (A). The control mice that did not receive amoxicillin had significantly more abundance of *Dubosiella* and *Lachnispiraceae NK4A136 group*. Volcano plot showing differential abundance of bacteria on Day 42 between the two groups (B). The amoxicillin exposed mice had higher abundance of *Desulfovibrio* and *Mucispirillum* but lower abundance of *Ureaplasma*. Day 42 biplots show that *Dubosiella*, *Lactobacillus,* and *Lachnospiraceae NK4A136 group* were three of the dominant species driving the changes in clustering (C).

**Supplementary Figure 2:**
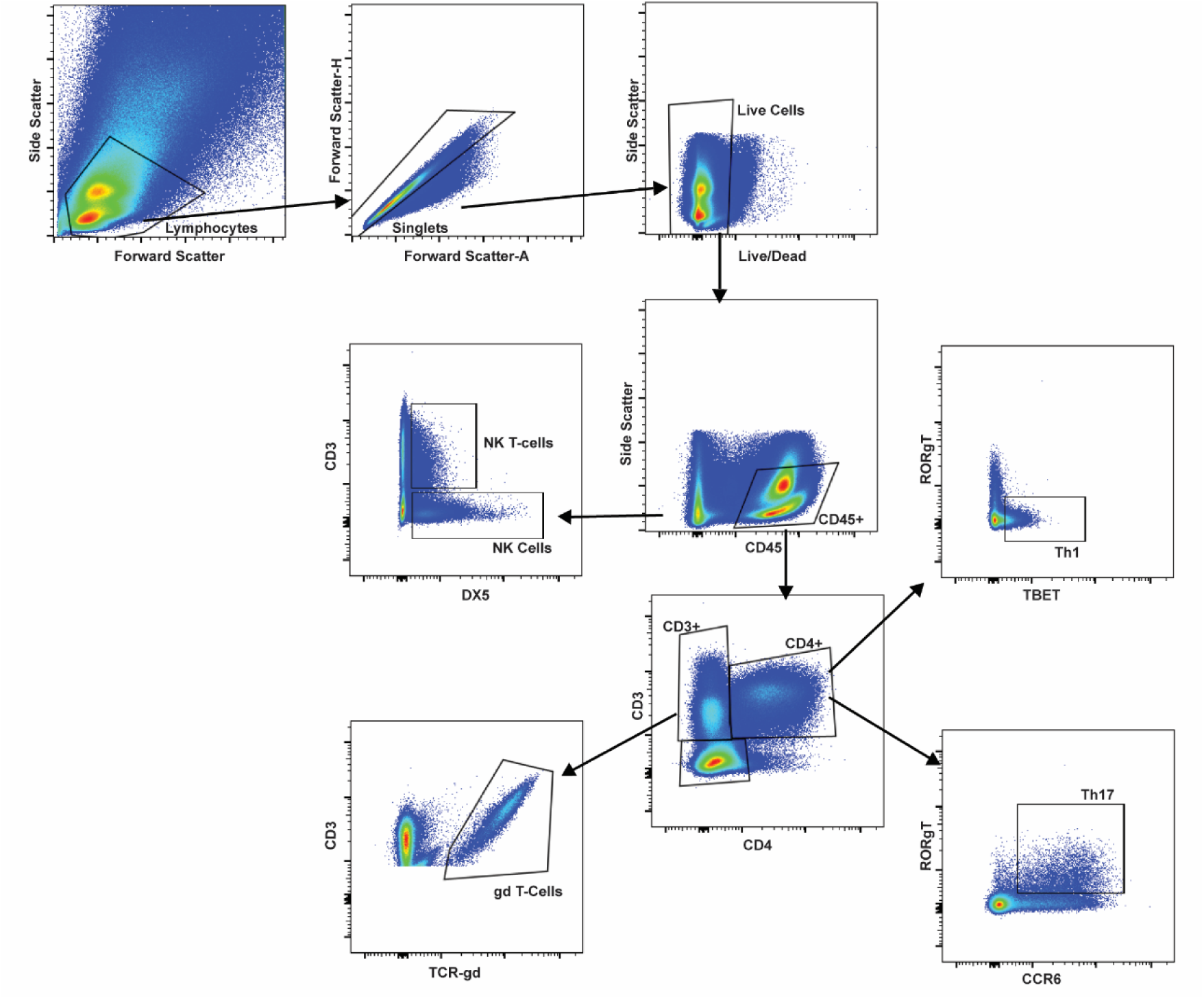
Flow cytometry gating strategy: Cells were initially selected based on their forward and side scatter characteristics. This cell population was further selected for single cells that were alive and then for the CD45+ expressing leukocytes. CD45+ cells were further gated either based on DX5 expression of CD3CD4 expression. NK cells were defined as CD45+DX5+CD3-, NK T-cells were defined as CD45+DX5+CD3+. CD45+CD3+CD4- cells were further gated on TCRgd expression. γδ T-cells were defined as CD45+CD3+CD4-TCRgd+ cells. The CD3+CD4+ double positive population was then gated on CCR6 and RORγT expression. Th17 cells were defined as CD45+CD3+CD4+CCR6+RORγT+ cells. Th1 cells were defined as CD45+CD3+CD4+TBET+RORγT-.

**Supplementary Figure 3:**
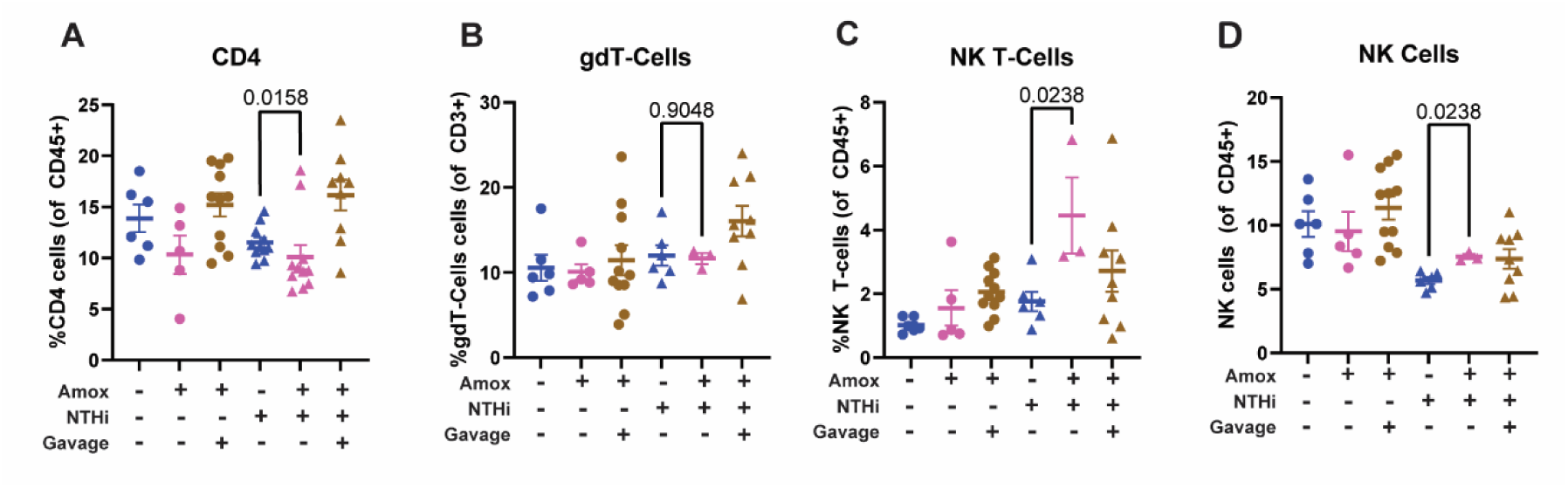
Amoxicillin effects on different lymphocyte populations in the lungs: Three weeks after antibiotic cessation, there was a significant decrease observed in the proportion of CD4+ CD45+ cells compared to the untreated controls, and this was restored with microbiome transfer (A). There was no difference between percentages of γδ T-cells in any of the conditions tested (B). There appeared to be an increase in NK T (C) and NK cells (D) in the amoxicillin treated group. Significance between amoxicillin treated and untreated mice in the NTHi exposed group was determined using a Mann-Whitney test. N=3-11, 1-3 replicates performed per condition.

**Supplementary Figure 4:**
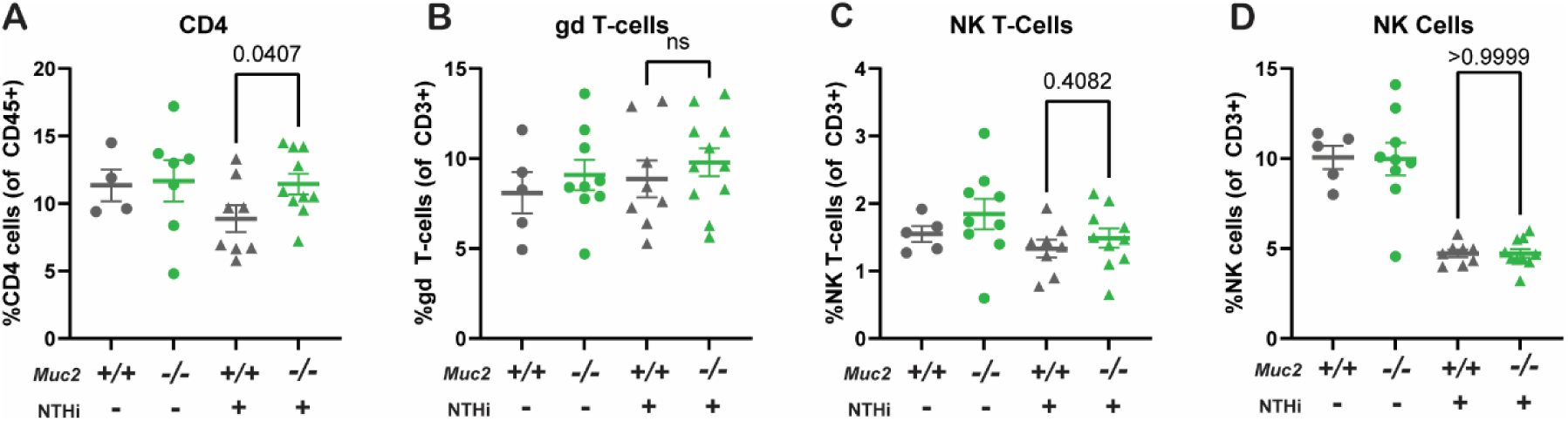
Gut mucin composition effects on different lymphocyte populations in the lungs: *Muc2^+/+^* and *Muc2^-/-^* mice were assessed for different lymphocyte populations by flow cytometry in the presence and absence of sterile inflammation (aerosolized NTHi). In the presence of NTHi, *Muc2^-/^* mice had increased CD4+ CD45+ cells compared to the control animals (A). There was no difference in percentages of γδ T-cells (B), NK T-cells (C), or NK Cells (D) in any of the conditions tested between the *Muc2^+/+^* and *Muc2^-/-^* mice. Significance between *Muc2^+/+^*and *Muc2^-/-^* mice in the NTHi exposed group was determined using a Mann-Whitney test. N=5-10, 2 replicates per condition were performed.

**Supplementary Figure 5:**
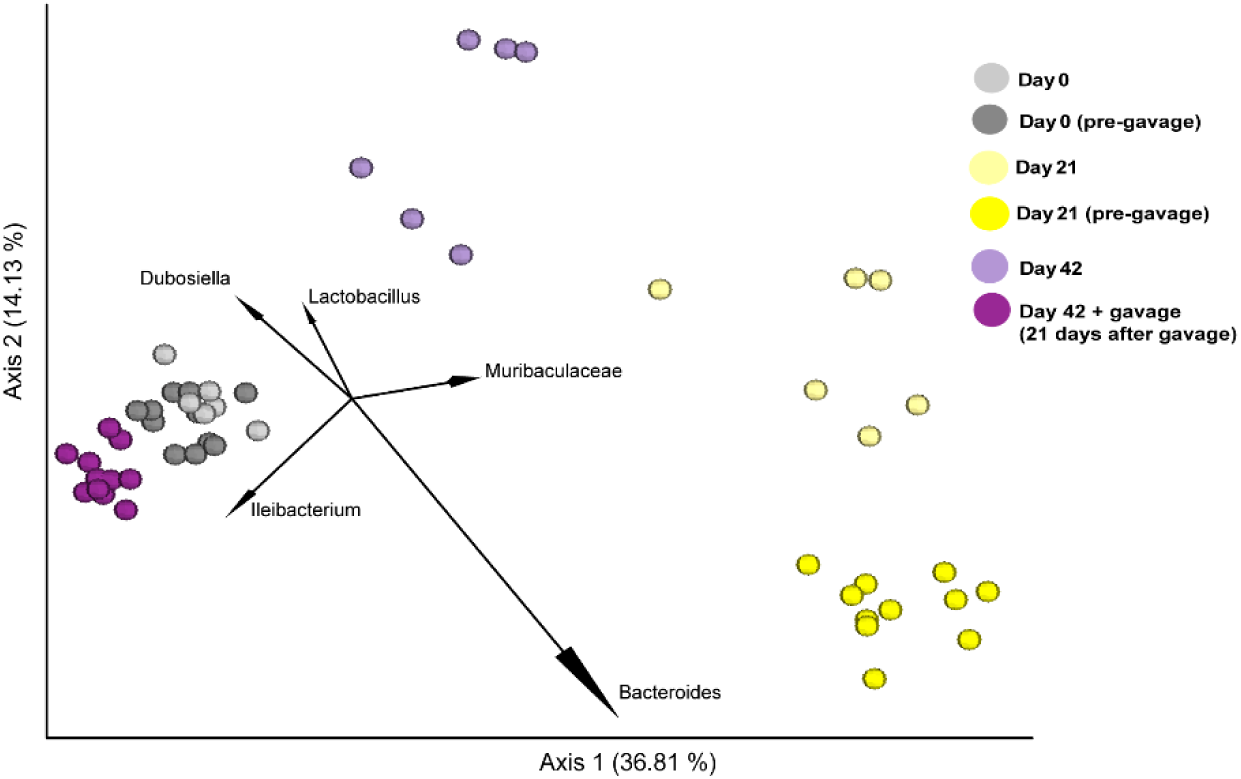
Fecal reconstitution restores beta-diversity in amoxicillin treated mice. Beta-diversity PCA plot shows the different bacterial species that were important for independent clustering.

**Supplementary Figure 6:**
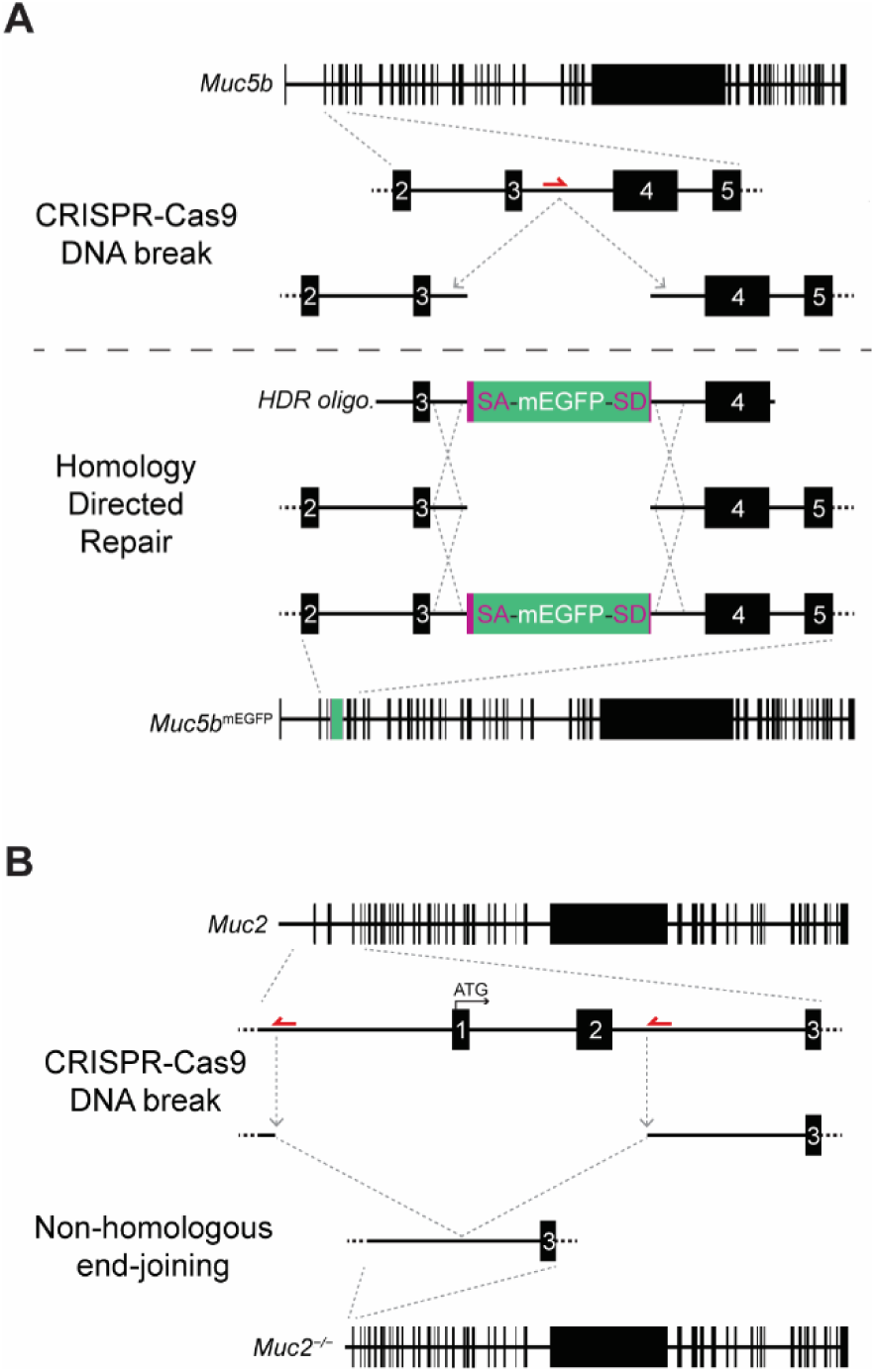
Maps for Genetically Engineered Mouse Strains. **A.** *Muc5b*^mEGFP^ knock-in mice were produced by insertion of a splice acceptor (SA)-mEGFP-splice donor (SD) construct that encoded a synthetic in-from exon between endogenous exons 3 and 4. **B.** *Muc2* knockout mice were produced by targeting with gRNAs that produced DNA breaks in the *Muc2* 5-flanking region and intron 2. This results in loss of the core Muc2 promoter and the introduction of frameshift mutations from remaining nearby open reading frames.

## References

1. Bradley, C.P., Teng, F., Felix, K.M., Sano, T., Naskar, D., Block, K.E., Huang, H., Knox, K.S., Littman, D.R., and Wu, H.-J.J. (2017). Segmented Filamentous Bacteria Provoke Lung Autoimmunity by Inducing Gut-Lung Axis Th17 Cells Expressing Dual TCRs. Cell Host & Microbe 22, 697–704.e4. 10.1016/j.chom.2017.10.007.

2. Jin, S., Zhao, D., Cai, C., Song, D., Shen, J., Xu, A., Qiao, Y., Ran, Z., and Zheng, Q. (2017). Low-dose penicillin exposure in early life decreases Th17 and the susceptibility to DSS colitis in mice through gut microbiota modification. Sci Rep 7, 43662. 10.1038/srep43662.

3. Goto, Y., Panea, C., Nakato, G., Cebula, A., Lee, C., Diez, M.G., Laufer, T.M., Ignatowicz, L., and Ivanov, I.I. (2014). Segmented Filamentous Bacteria Antigens Presented by Intestinal Dendritic Cells Drive Mucosal Th17 Cell Differentiation. Immunity 40, 594–607. 10.1016/j.immuni.2014.03.005.

4. Chen, B., Chen, H., Shu, X., Yin, Y., Li, J., Qin, J., Chen, L., Peng, K., Xu, F., Gu, W., et al. (2018). Presence of Segmented Filamentous Bacteria in Human Children and Its Potential Role in the Modulation of Human Gut Immunity. Frontiers in Microbiology 9.

5. Sun, C.-Y., Yang, N., Zheng, Z.-L., Liu, D., and Xu, Q.-L. (2023). T helper 17 (Th17) cell responses to the gut microbiota in human diseases. Biomed Pharmacother 161, 114483. 10.1016/j.biopha.2023.114483.

6. Chen, K., and Kolls, J.K. (2013). T cell-mediated host immune defenses in the lung. Annu Rev Immunol 31, 605–633. 10.1146/annurev-immunol-032712-100019.

7. Dubin, P.J., and Kolls, J.K. (2008). Th17 cytokines and mucosal immunity. Immunological Reviews 226, 160–171. 10.1111/j.1600-065X.2008.00703.x.

8. Yasuda, K., Takeuchi, Y., and Hirota, K. (2019). The pathogenicity of Th17 cells in autoimmune diseases. Semin Immunopathol 41, 283–297. 10.1007/s00281-019-00733-8.

9. Brown, R.L., Sequeira, R.P., and Clarke, T.B. (2017). The microbiota protects against respiratory infection via GM-CSF signaling. Nat Commun 8, 1512. 10.1038/s41467-017-01803-x.

10. Kudva, A., Scheller, E.V., Robinson, K.M., Crowe, C.R., Choi, S.M., Slight, S.R., Khader, S.A., Dubin, P.J., Enelow, R.I., Kolls, J.K., et al. (2011). Influenza A inhibits Th17-mediated host defense against bacterial pneumonia in mice. J. Immunol. 186, 1666–1674. 10.4049/jimmunol.1002194.

11. Orlov, M., Dmyterko, V., Wurfel, M.M., and Mikacenic, C. (2017). Th17 cells are associated with protection from ventilator associated pneumonia. PLoS ONE 12, e0182966. 10.1371/journal.pone.0182966.

12. Prescott, H.C., Langa, K.M., and Iwashyna, T.J. (2015). Readmission diagnoses after hospitalization for severe sepsis and other acute medical conditions. JAMA 313, 1055–1057. 10.1001/jama.2015.1410.

13. Baggs, J., Jernigan, J.A., Halpin, A.L., Epstein, L., Hatfield, K.M., and McDonald, L.C. (2018). Risk of Subsequent Sepsis Within 90 Days After a Hospital Stay by Type of Antibiotic Exposure. Clinical Infectious Diseases 66, 1004–1012. 10.1093/cid/cix947.

14. Yang, Y., Torchinsky, M.B., Gobert, M., Xiong, H., Xu, M., Linehan, J.L., Alonzo, F., Ng, C., Chen, A., Lin, X., et al. (2014). Focused specificity of intestinal TH17 cells towards commensal bacterial antigens. Nature 510, 152–156. 10.1038/nature13279.

15. Yin, Y., Wang, Y., Zhu, L., Liu, W., Liao, N., Jiang, M., Zhu, B., Yu, H.D., Xiang, C., and Wang, X. (2013). Comparative analysis of the distribution of segmented filamentous bacteria in humans, mice and chickens. ISME J 7, 615–621. 10.1038/ismej.2012.128.

16. Ivanov, I.I., Zhou, L., and Littman, D.R. (2007). Transcriptional regulation of Th17 cell differentiation. Seminars in Immunology 19, 409–417. 10.1016/j.smim.2007.10.011.

17. Dang, A.T., and Marsland, B.J. (2019). Microbes, metabolites, and the gut–lung axis. Mucosal Immunology 12, 843–850. 10.1038/s41385-019-0160-6.

18. Huang, Y., Mao, K., Chen, X., Sun, M.-A., Kawabe, T., Li, W., Usher, N., Zhu, J., Urban, J.F., Paul, W.E., et al. (2018). S1P-dependent interorgan trafficking of group 2 innate lymphoid cells supports host defense. Science 359, 114–119. 10.1126/science.aam5809.

19. Fan, T., Tai, C., Sleiman, K.C., Cutcliffe, M.P., Kim, H., Liu, Y., Li, J., Xin, G., Grashel, M., Baert, L., et al. (2025). Aberrant T follicular helper cells generated by TH17 cell plasticity in the gut promote extraintestinal autoimmunity. Nat Immunol 26, 790–804. 10.1038/s41590-025-02125-7.

20. Ni, S., Yuan, X., Cao, Q., Chen, Y., Peng, X., Lin, J., Li, Y., Ma, W., Gao, S., and Chen, D. (2023). Gut microbiota regulate migration of lymphocytes from gut to lung. Microbial Pathogenesis 183, 106311. 10.1016/j.micpath.2023.106311.

21. Armstrong, G., Cantrell, K., Huang, S., McDonald, D., Haiminen, N., Carrieri, A.P., Zhu, Q., Gonzalez, A., McGrath, I., Beck, K.L., et al. (2021). Efficient computation of Faith’s phylogenetic diversity with applications in characterizing microbiomes. Genome Res. 31, 2131–2137. 10.1101/gr.275777.121.

22. Faith, D.P. (1992). Conservation evaluation and phylogenetic diversity. Biological Conservation 61, 1– 10. 10.1016/0006-3207(92)91201-3.

23. Pielou, E.C. (1966). The measurement of diversity in different types of biological collections. Journal of Theoretical Biology 13, 131–144. 10.1016/0022-5193(66)90013-0.

24. Lozupone, C., and Knight, R. (2005). UniFrac: a new phylogenetic method for comparing microbial communities. Appl Environ Microbiol 71, 8228–8235. 10.1128/AEM.71.12.8228-8235.2005.

25. Bolyen, E., Rideout, J.R., Dillon, M.R., Bokulich, N.A., Abnet, C.C., Al-Ghalith, G.A., Alexander, H., Alm, E.J., Arumugam, M., Asnicar, F., et al. (2019). Reproducible, interactive, scalable and extensible microbiome data science using QIIME 2. Nat Biotechnol 37, 852–857. 10.1038/s41587-019-0209-9.

26. Mandal, S., Van Treuren, W., White, R.A., Eggesbø, M., Knight, R., and Peddada, S.D. (2015). Analysis of composition of microbiomes: a novel method for studying microbial composition. Microb Ecol Health Dis 26, 27663. 10.3402/mehd.v26.27663.

27. Moghaddam, S.J., Clement, C.G., De la Garza, M.M., Zou, X., Travis, E.L., Young, H.W.J., Evans, C.M., Tuvim, M.J., and Dickey, B.F. (2008). Haemophilus influenzae lysate induces aspects of the chronic obstructive pulmonary disease phenotype. Am J Respir Cell Mol Biol 38, 629–638. 10.1165/rcmb.2007-0366OC.

28. Li, W., Zhang, X., Yang, Y., Yin, Q., Wang, Y., Li, Y., Wang, C., Wong, S.M., Wang, Y., Goldfine, H., et al. (2018). Recognition of conserved antigens by Th17 cells provides broad protection against pulmonary Haemophilus influenzae infection. Proc Natl Acad Sci U S A 115, E7149–E7157. 10.1073/pnas.1802261115.

29. Wenzel, U.A., Jonstrand, C., Hansson, G.C., and Wick, M.J. (2015). CD103+CD11b+ Dendritic Cells Induce Th17 T Cells in Muc2-Deficient Mice with Extensively Spread Colitis. PLoS One 10, e0130750. 10.1371/journal.pone.0130750.

30. Shan, M., Gentile, M., Yeiser, J.R., Walland, A.C., Bornstein, V.U., Chen, K., He, B., Cassis, L., Bigas, A., Cols, M., et al. (2013). Mucus Enhances Gut Homeostasis and Oral Tolerance by Delivering Immunoregulatory Signals. Science 342, 447–453. 10.1126/science.1237910.

31. Van der Sluis, M., De Koning, B.A.E., De Bruijn, A.C.J.M., Velcich, A., Meijerink, J.P.P., Van Goudoever, J.B., Büller, H.A., Dekker, J., Van Seuningen, I., Renes, I.B., et al. (2006). Muc2-deficient mice spontaneously develop colitis, indicating that MUC2 is critical for colonic protection. Gastroenterology 131, 117–129. 10.1053/j.gastro.2006.04.020.

32. Yoon, S., Lee, G., Yu, J., Lee, K., Lee, K., Si, J., You, H.J., and Ko, G. (2022). Distinct Changes in Microbiota-Mediated Intestinal Metabolites and Immune Responses Induced by Different Antibiotics. Antibiotics (Basel) 11, 1762. 10.3390/antibiotics11121762.

33. The Most Common Antibiotics Prescribed in 2023 https://www.definitivehc.com/resources/healthcare-insights/most-prescribed-antibiotics.

34. Lalmohamed, A., Venekamp, R.P., Bolhuis, A., Souverein, P.C., van de Wijgert, J.H.H.M., Gulliford, M.C., and Hay, A.D. (2024). Within-episode repeat antibiotic prescriptions in patients with respiratory tract infections: A population-based cohort study. Journal of Infection 88, 106135. 10.1016/j.jinf.2024.106135.

35. Tirelle, P., Breton, J., Riou, G., Déchelotte, P., Coëffier, M., and Ribet, D. (2020). Comparison of different modes of antibiotic delivery on gut microbiota depletion efficiency and body composition in mouse. BMC Microbiol 20, 340. 10.1186/s12866-020-02018-9.

36. Livanos, A.E., Greiner, T.U., Vangay, P., Pathmasiri, W., Stewart, D., McRitchie, S., Li, H., Chung, J., Sohn, J., Kim, S., et al. (2016). Antibiotic-mediated gut microbiome perturbation accelerates development of type 1 diabetes in mice. Nat Microbiol 1, 16140. 10.1038/nmicrobiol.2016.140.

37. Netea, M.G., Joosten, L.A.B., Latz, E., Mills, K.H.G., Natoli, G., Stunnenberg, H.G., O’Neill, L.A.J., and Xavier, R.J. (2016). Trained immunity: a program of innate immune memory in health and disease. Science 352, aaf1098. 10.1126/science.aaf1098.

38. Doan, T.A., Forward, T.S., Schafer, J.B., Lucas, E.D., Fleming, I., Uecker-Martin, A., Ayala, E., Guthmiller, J.J., Hesselberth, J.R., Morrison, T.E., et al. (2024). Immunization-induced antigen archiving enhances local memory CD8+ T cell responses following an unrelated viral infection. npj Vaccines 9, 66. 10.1038/s41541-024-00856-6.

39. Anthony, W.E., Wang, B., Sukhum, K.V., D’Souza, A.W., Hink, T., Cass, C., Seiler, S., Reske, K.A., Coon, C., Dubberke, E.R., et al. (2022). Acute and persistent effects of commonly used antibiotics on the gut microbiome and resistome in healthy adults. Cell Rep 39, 110649. 10.1016/j.celrep.2022.110649.

40. Palleja, A., Mikkelsen, K.H., Forslund, S.K., Kashani, A., Allin, K.H., Nielsen, T., Hansen, T.H., Liang, S., Feng, Q., Zhang, C., et al. (2018). Recovery of gut microbiota of healthy adults following antibiotic exposure. Nat Microbiol 3, 1255–1265. 10.1038/s41564-018-0257-9.

41. Zhang, Y., Tu, S., Ji, X., Wu, J., Meng, J., Gao, J., Shao, X., Shi, S., Wang, G., Qiu, J., et al. (2024). Dubosiella newyorkensis modulates immune tolerance in colitis via the L-lysine-activated AhR-IDO1-Kyn pathway. Nat Commun 15, 1333. 10.1038/s41467-024-45636-x.

42. Luo, Z., Liao, G., Meng, M., Huang, X., Liu, X., Wen, W., Yue, T., Yu, W., Wang, C., and Jiang, Y. (2025). The Causal Relationship Between Gut and Skin Microbiota and Chronic Obstructive Pulmonary Disease:A Bidirectional Two-Sample Mendelian Randomization Analysis. Int J Chron Obstruct Pulmon Dis 20, 709–722. 10.2147/COPD.S494289.

43. Zhang, M., Liang, Y., Liu, Y., Li, Y., Shen, L., and Shi, G. (2022). High-fat diet-induced intestinal dysbiosis is associated with the exacerbation of Sjogren’s syndrome. Front Microbiol 13, 916089. 10.3389/fmicb.2022.916089.

44. Pemmasani, G., Loftus, E.V., and Tremaine, W.J. (2022). Prevalence of Pulmonary Diseases in Association with Inflammatory Bowel Disease. Dig Dis Sci 67, 5187–5194. 10.1007/s10620-022-07385-z.

45. Wu, M., Wu, Y., Li, J., Bao, Y., Guo, Y., and Yang, W. (2018). The Dynamic Changes of Gut Microbiota in Muc2 Deficient Mice. International Journal of Molecular Sciences 19, 2809. 10.3390/ijms19092809.

46. Paudel, K.R., Dharwal, V., Patel, V.K., Galvao, I., Wadhwa, R., Malyla, V., Shen, S.S., Budden, K.F., Hansbro, N.G., Vaughan, A., et al. (2020). Role of Lung Microbiome in Innate Immune Response Associated With Chronic Lung Diseases. Front. Med. 7. 10.3389/fmed.2020.00554.

47. Zhu, Y., Chen, B., Zhang, X., Akbar, M.T., Wu, T., Zhang, Y., Zhi, L., and Shen, Q. (2024). Exploration of the Muribaculaceae Family in the Gut Microbiota: Diversity, Metabolism, and Function. Nutrients 16. 10.3390/nu16162660.

48. Wu, M., Wu, Y., Li, J., Bao, Y., Guo, Y., and Yang, W. (2018). The Dynamic Changes of Gut Microbiota in Muc2 Deficient Mice. International Journal of Molecular Sciences 19, 2809. 10.3390/ijms19092809.

49. Spivak, I., Fluhr, L., and Elinav, E. (2022). Local and systemic effects of microbiome-derived metabolites. EMBO Rep 23, e55664. 10.15252/embr.202255664.

50. Ahmed, H., Leyrolle, Q., Koistinen, V., Kärkkäinen, O., Layé, S., Delzenne, N., and Hanhineva, K. (2022). Microbiota-derived metabolites as drivers of gut-brain communication. Gut Microbes 14, 2102878. 10.1080/19490976.2022.2102878.

51. Fakih, D., Rodriguez-Piñeiro, A.M., Trillo-Muyo, S., Evans, C.M., Ermund, A., and Hansson, G.C. (2020). Normal murine respiratory tract has its mucus concentrated in clouds based on the Muc5b mucin. Am J Physiol Lung Cell Mol Physiol 318, L1270–L1279. 10.1152/ajplung.00485.2019.

52. Velcich, A., Yang, W., Heyer, J., Fragale, A., Nicholas, C., Viani, S., Kucherlapati, R., Lipkin, M., Yang, K., and Augenlicht, L. (2002). Colorectal cancer in mice genetically deficient in the mucin Muc2. Science 295, 1726–1729. 10.1126/science.1069094.

53. Van der Sluis, M., De Koning, B.A.E., De Bruijn, A.C.J.M., Velcich, A., Meijerink, J.P.P., Van Goudoever, J.B., Büller, H.A., Dekker, J., Van Seuningen, I., Renes, I.B., et al. (2006). Muc2-deficient mice spontaneously develop colitis, indicating that MUC2 is critical for colonic protection. Gastroenterology 131, 117–129. 10.1053/j.gastro.2006.04.020.

54. Thompson, L.R., Sanders, J.G., McDonald, D., Amir, A., Ladau, J., Locey, K.J., Prill, R.J., Tripathi, A., Gibbons, S.M., Ackermann, G., et al. (2017). A communal catalogue reveals Earth’s multiscale microbial diversity. Nature 551, 457–463. 10.1038/nature24621.

55. 16S Illumina Amplicon Protocol : earthmicrobiome https://earthmicrobiome.ucsd.edu/protocols-and-standards/16s/.

56. Callahan, B.J., McMurdie, P.J., Rosen, M.J., Han, A.W., Johnson, A.J.A., and Holmes, S.P. (2016). DADA2: High-resolution sample inference from Illumina amplicon data. Nat Methods 13, 581–583. 10.1038/nmeth.3869.

57. Janssen, S., McDonald, D., Gonzalez, A., Navas-Molina, J.A., Jiang, L., Xu, Z.Z., Winker, K., Kado, D.M., Orwoll, E., Manary, M., et al. (2018). Phylogenetic Placement of Exact Amplicon Sequences Improves Associations with Clinical Information. mSystems 3, 10.1128/msystems.00021-18. 10.1128/msystems.00021-18.

58. Vázquez-Baeza, Y., Pirrung, M., Gonzalez, A., and Knight, R. (2013). EMPeror: a tool for visualizing high-throughput microbial community data. Gigascience 2, 2047–217X-2–16. 10.1186/2047-217X-2-16.

